# orthogene: a Bioconductor package to easily map genes within and across hundreds of species

**DOI:** 10.64898/2026.01.17.700094

**Authors:** Brian M. Schilder, Alan E. Murphy, Nathan G. Skene

## Abstract

**Motivation:** Mapping genes across identifier systems and species is a routine but critical step in bioinformatics workflows. Despite its ubiquity, gene mapping is frequently handled using bespoke, ad hoc solutions, increasing duplicated effort and introducing opportunities for error. These issues are exacerbated by the prevalence of non–one-to-one homolog relationships and inconsistent handling of gene identifiers across species and databases, which can compromise downstream analyses and reproducibility.

**Results:** We present `orthogene`, an R/Bioconductor package that simplifies gene mapping within and across hundreds of species. `orthogene` provides a unified, workflow-oriented framework that integrates automated species and identifier standardization, homolog inference across multiple databases, flexible handling of ambiguous homolog relationships, and transformation of gene lists, tables, and high-dimensional matrices into analysis-ready formats. By abstracting common sources of technical complexity while retaining user control, `orthogene` enables transparent, reproducible, and scalable gene mapping across a wide range of biological contexts.

**Availability:** https://bioconductor.org/packages/orthogene

## 1 Introduction

Mapping genes from one ID system to another, whether within or across across species, is a near-ubiquitous task in bioinformatics. Despite this, many researchers find themselves independently reinventing their own approaches to accomplish this objective, leading to massively duplicated efforts. Furthermore, given how far upstream gene mapping is to many bioinformatics pipelines, suboptimal solutions (e.g. incorrect mappings or genes being dropped entirely) can greatly impact downstream analyses and conclusions^1^. Complicating this matter further is the fact that many gene mappings are inherently not one to one, such as when there are multiple homologs of the same gene in one species but only one version in another species^2^. Arbitrary and often undocumented, choices in how to handle these situations can inadvertently contribute to the scientific reproducibility crisis^3,4^. Here we introduce `orthogene`, a Bioconductor R package that drastically simplifies intra- and inter-species gene mapping.

By integrating a comprehensive suite of gene mapping databases alongside fully automated strategies to handle homology mapping and filtering, `orthogene` abstracts away some of the common and tedious tasks in bioinformatics while still providing users with extensive control if needed. Presently, it is able to map genes within and across 760 different organisms spanning the tree of life. This obviates the need for researchers to independently develop unique solutions to map between each ID coordinate system (Ensembl transcripts vs. Ensembl genes vs. HGNC gene symbols vs. Entrez vs. UniProt, and all combinations thereof) or between each pair of species (mouse to human, human to mouse, mouse to nematode, fly to zebrafish, etc.). Instead, `orthogene` offers a unified programmatic solution to address this wide range of scenarios. Furthermore, it employs different routines to automatically infer, ingest, and process a wide variety of data input formats (various classes of lists, tables, matrices) and provides output data in whichever format best meets the user’s needs.

Through automation, robust and flexible handling of homolog ambiguity, and immediate usability, we believe that `orthogene` will serve as a tool to accelerate research across the life sciences.

## 2 Methods

### 2.1 Implementation and installation

`orthogene` is implemented in R and distributed via Bioconductor, ensuring standardized release cycles, dependency management, and long-term maintenance. It can also be installed from source via GitHub. Moreover we provide a fully containerized Docker environment with all system and R dependencies preinstalled, further enhancing usability and reproducibility via a single line of code (docker pull ghcr.io/neurogenomics/orthogene). Multiple layers of Continuous Integration ensure robust testing on both GitHub (using `rworkflows`^5^) and the Bioconductor service^6^, which provide automated code checks, code coverage reports, automated documentation website updates, and automated updates to the official Docker container distribution. Before each new version deployment, `orthogene` is extensively and regularly pressure-tested across multiple operating systems (Linux, Mac, Windows), achieving a code coverage score of *>*84% (https://app.codecov.io/gh/neurogenomics/orthogene). All source code is openly available under a permissive GPL-3 license and distributed on GitHub (https://github.com/neurogenomics/orthogene). `orthogene` was first released under Bioc version 3.14 (requires R version ≥ 4.1) and is available in all subsequent releases of Bioc; presently version 3.22 (https://bioconductor.org/packages/release/bioc/html/orthogene.html). All functions and datasets are fully documented within the package, as well as on a dedicated website, which also includes tutorials (https://neurogenomics.github.io/orthogene/).

### 2.2 Input standardization

Designed to be flexible and highly automated, `orthogene`’s primary high-level function `convert_orthologs()` can take a variety of input data classes (`list`, `vector`, `data.frame`, `data.table`, `tibble`, `matrix`, `sparseMatrix`, or `DelayedArray`^7^) and infers how to appropriately process the data, unless otherwise specified by the user (Fig. 1a). First, it maps the user-provided species name onto a standardized nomenclature, e.g. “human” to “Homo sapiens” (Fig. 1b, *top*). If the species is unknown, as can sometimes happen when data is poorly documented, `infer_species()` identifies the most likely species by computing the percent of matching gene IDs against a variety of reference organisms (see vignette demonstrating the reliability of this method https://neurogenomics.github.io/orthogene/articles/infer_species.html). Optionally, the genes can also be mapped onto a standardized gene nomenclature (Fig. 1b, *bottom*) by leveraging the `g:Profiler` Application Programming Interface (API)^8,9^ which in turn queries a variety of databases like NCBI (National Center for Biotechnology Information)^10^ and Ensembl^11^.

**Figure 1.**
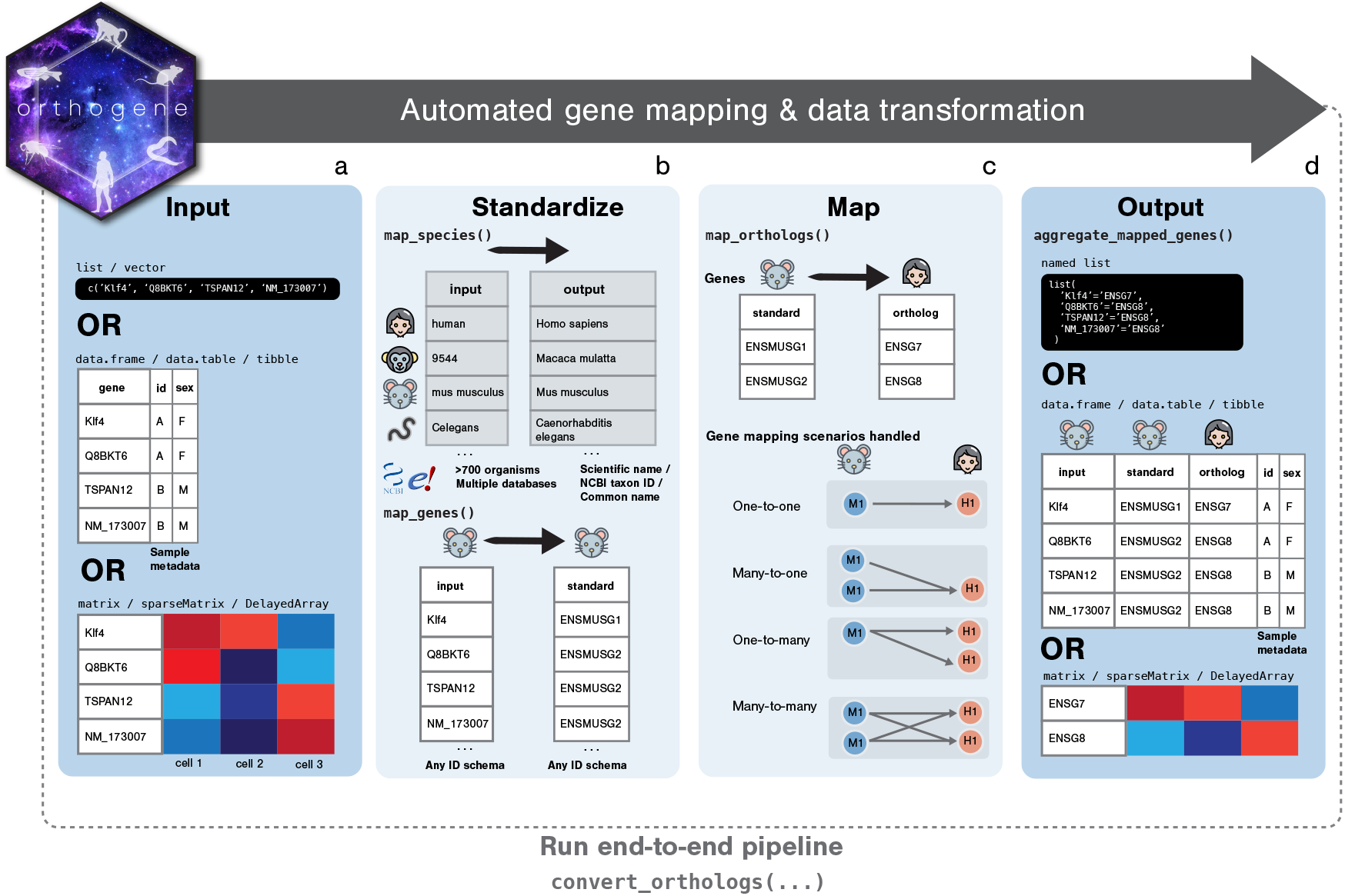
Overview of the orthogene workflow. The homolog mapping pipeline can be run with a unified wrapper function (`convert_orthologs`). Alternatively, individual subfunctions can be called directly by users who require more precise control over each step (**a-d**). **a**, `orthogene` can flexibly take various input data classes and automatically infer how to process them. **b**, The `map_species()` function standardizes species names onto a common nomenclature, such as scientific names (e.g. *Homo sapiens*), NCBI taxon IDs (e.g. 9606), or common names (e.g. human). Optionally, `map_genes()` harmonizes all input genes onto a common standard, which is especially helpful when using a mix of ID systems. **c**, Genes are then translated across species using one of several backends (`gprofiler2`, `homologene`, or `babelgene`). **d**, Finally, mapped genes can be returned in different formats, including a table (*top*) a dictionary (*middle*), or an aggregated matrix (*bottom*) using the `aggregate_mapped_genes()` function.

### 2.3 Cross-species mapping

Next, homologous genes are mapped across species (Fig. 1c, *top*) using one of several optional backends (`gprofiler2`^9^, `homologene`^12^, `babelgene`^13^). `orthogene` supports multiple homology inference backends to accommodate different analytical requirements, including both remote, broad-coverage services (`gprofiler2`) and fast local resources (`homologene`, `babelgene`). In addition to performance differences, use of local resources enables `orthogene` to operate in compute environments where stable internet access is limited due to technical or security constraints. A detailed comparison of backend strengths and limitations is provided in Table S1.

A key feature distinguishing `orthogene` from other inter-species mapping tools is its support for multiple strategies (Table S2) to handle both orthologous (one-to-one) and paralogous (many-to-one, one-to-many, or many-to-many) relationships (Fig. 1c, *bottom*). Strategies can be easily switched using the `non121_strategy` argument. Most existing R packages focus solely on returning unfiltered mappings. A partial exception is `babelgene`, which by default selects a single homolog per input based on an evidence-ranking heuristic (also implemented in `orthogene` as the “*keep_popular*” strategy).

By default, `orthogene` applies the “*drop_both_species*” strategy, which removes genes with multiple cross-species mappings, retaining only one-to-one orthologs as is common practice in bioinformatics. This approach is often justified by the assumption that non–one-to-one orthologs are more likely to have functionally diverged^2,14^, although competing evidence exists^1,15,16^. Importantly, `orthogene` enables this simple default while allowing users to systematically apply alternative strategies when appropriate. To facilitate rapid quality assessment, `orthogene` automatically generates reports summarizing the inferred species, number of unique input genes, conversion rates, and genes dropped at each filtering step. These reports allow users to quickly compare mapping strategies and quantify their effects on gene retention prior to downstream analyses.

### 2.4 Output transformation

Finally, the user’s data is returned in one of multiple formats, depending on the input data and the user-specified arguments. Results can be returned as a table that traces the relationships between inputs genes, intra-species mapped genes, and interspecies mapped genes (Fig. 1d, *top*). Alternatively, a dictionary (named list) of key:value (input:output) pairs can be returned. If the user provided a matrix with numerical data as input (with row or column names representing genes) a new transformed matrix will be returned. If genes were row names, the new matrix map will have fewer rows than the original input data due to dropped genes that did not meet the mapping criteria (e.g. one-to-one) or did not have any known homologs between the two species. In cases where multiple genes from the input species (e.g. mouse) map onto a single gene in the output species (e.g. human) or vice versa, the user can specify an aggregation/splitting function to transform the matrix accordingly (e.g. “*mean*”=take the average value when condensing gene vectors).

Beyond interspecies conversion, `orthogene` is also very helpful for aggregating data that has been mapped across gene ID systems within the same species. For example, one might have a single-cell gene expression matrix with each row labeled with Ensembl transcript IDs (ENST). `orthogene` makes the task of converting this matrix into a gene-level representation easy by mapping the transcript IDs to gene IDs (such as ENSG or HGNC) and then aggregating with a preferred method (mean, median, min, max, etc). To ensure these functions are scalable to large datasets, we have further optimized them with chunking operations^7^.

### 2.5 Additional utilities

In addition to these core functions, `orthogene` also contains a variety of utility function that can be helpful for common bioinformatics workflows. `all_species()`: Provides a table of all species available via each backend method. `all_genes()`: Provides a table of all genes from a particular species. `create_background()`: Generates a gene background as the union (or intersect) of all orthologs between an input species and the output species to be mapped to. This can be useful when generating random lists of background genes to test against in analyses with data from multiple species (e.g. enrichment of mouse cell-type markers gene sets in human population-derived gene sets). `get_silhouettes()`: Maps species names and downloads silhouettes of requested species from PhyloPic (phylopic.org/) via the R package `rphylopic`^17^. `plot_orthotree()`: Automatically creates a phylogenetic tree plot annotated with metadata describing how many orthologous genes each species shares with a given reference species (“human” by default; see Fig. S2 for example).

### 2.6 Community uptake

Despite having not yet been thoroughly advertised, `orthogene` has already seen considerable uptake by the R community. Since its initial release, `orthogene` has been installed from Bioconductor 26,435 times by 17,035 unique IP addresses, averaging approximately 755 downloads per month (see Fig. S3; data from: https://bioconductor.org/packages/stats/bioc/orthogene/). By downloads, it currently ranks 308/2361 (top 13% percentile) of all Bioconductor packages. Meanwhile, other R packages have already begun to build upon `orthogene` by including it as a dependency, such as `BulkSignalR`^18^ and `sparrow`^19^. Together, this suggests that the R community is already finding value in `orthogene`.

## 3 Benchmarks

Integrating multiple homology conversion backends (`gprofiler2`, `homologene`, `babelgene`) into a single tool enabled systematic benchmarking of coverage and robustness across methods. We therefore defined two benchmark tasks. First, we exhaustively retrieved all genome-wide genes from 20 different model organisms using the `all_genes()` function. Second, we converted each organism’s genes to human homologs using `convert_orthologs()`, with the goal of maximizing recovery of one-to-one homologs. As a control to ensure genes were not lost due to preprocessing, we also converted all human genes to themselves following gene nomenclature standardization. For both tasks, we recorded runtime per backend, as performance may be an important consideration for some users.

In the `all_genes()` task, `gprofiler2` consistently retrieved more unique genes than the other methods for vertebrates (Fig. S4a), particularly in humans (2.1 × greater coverage) and mice (3.8 × greater coverage). This trend reversed for most invertebrates, plants, and microorganisms, reflecting the absence of these species from `gprofiler2`. On average, `homologene` was the fastest method (0.018 s/species), followed by `babelgene` (1.03 s/species) and `gprofiler2` (3.7 s/species) (Fig. S4b). Notably, `gprofiler2` performed well despite relying on remote queries rather than local storage, aided in part by chunking optimizations implemented in `orthogene`.

In the `convert_orthologs()` task, all three methods performed comparably for vertebrates (Fig. S4c). However, `gprofiler2` consistently underperformed for non-vertebrate organisms, indicating less extensive coverage for these human– nonhuman pairs. As before, `gprofiler2` was slower (7.9 s/species) than `homologene` (0.13 s/species) and `babelgene` (0.09 s/species) (Fig. S4d).

Together, these results highlight a key strength of `orthogene`: a unified API that enables users to flexibly leverage complementary homology databases rather than relying on a single backend.

## 4 Discussion

Homologous gene relationships form the basis by which biological insight is transferred across species and are foundational to comparative and translational genomics. Analyses ranging from interpretation of perturbation experiments in model organisms to cross-species comparisons of transcriptional programs, pathway conservation, and disease gene prioritization all depend on reliable interspecies gene mapping. Despite this central role, homolog mapping is frequently treated as a preliminary preprocessing step rather than as a core analytical operation.

Several high-quality resources exist for homolog inference^**?**,8–12^. However, practical use typically requires additional steps to standardize species nomenclature, harmonize gene identifiers, select among competing homolog definitions, and integrate mappings into matrices, enrichment analyses, and visualizations. These steps are often implemented through bespoke scripts, reducing reproducibility, increasing technical burden, and introducing opportunities for subtle errors that may propagate through downstream analyses.

`orthogene` was developed to directly address these challenges through highly automated species and identifier standardization, homolog inference across multiple backends, explicit representation of complex homolog relationships, matrix transformation, background construction, and evolutionary contextualization within a unified and reproducible workflow. This design allows homolog mappings to be applied consistently across heterogeneous data types while preserving transparency and user control over methodological choices.

A central contribution of `orthogene` is its emphasis on analysis-ready representations. Direct transformation of high-dimensional gene-by-sample matrices using explicit homolog tables reduces reliance on ad hoc data manipulation and mitigates common mapping and duplication errors. Configurable aggregation strategies enable principled handling of one-to-many and many-to-many homolog relationships without mandating premature simplification that may distort biological signal.

Beyond mapping itself, `orthogene` addresses downstream statistical considerations that are frequently overlooked in cross-species analyses. In particular, construction of background gene universes that are consistent with species, homolog inference backend, and mapping strategy reduces bias in enrichment and overrepresentation tests. We make extensive use of this in our other software packages such as `EWCE`^20^ and `MSTExplorer`^21^.

Rather than replacing existing homolog databases, `orthogene` builds upon them by providing a consistent interface that renders their outputs directly usable within analytical workflows. Its sustained adoption within the Bioconductor ecosystem reflects the need for robust, workflow-integrated approaches to homolog-based analysis. Together, these features position `orthogene` as a practical and evolution-aware framework for reproducible cross-species genomics in R.

## 5 Data Availability

`orthogene` supporting data files: https://github.com/neurogenomics/orthogene/releases

## 6 Code Availability

`orthogene` GitHub: https://github.com/neurogenomics/orthogene

`gprofiler2` GitHub: https://github.com/cran/gprofiler2

`homologene` GitHub: https://github.com/oganm/homologene

`babelgene` GitHub: https://github.com/igordot/babelgene

## 7 Acknowledgments

This work was supported by a UK Dementia Research Institute (UK DRI) Future Leaders Fellowship [MR/T04327X/1] and the UK DRI which receives its funding from UK DRI Ltd, funded by the UK Medical Research Council, Alzheimer’s Society and Alzheimer’s Research UK.

We would like to thank the developers of `g:Profiler`/`gprofiler2`, `babelgene`, and `homologene` as well as all `orthogene` users for their invaluable support and feedback.

## 8 Author Contributions

BMS conceived of, designed, coded and wrote the manuscript for `orthogene`, with contributions from AEM. `orthogene`’s predecessor, `One2One`^22^, was developed by NGS.

