## Supplementary Materials for "orthogene: a Bioconductor package to easily map genes within and across hundreds of species"

### 1 Supplementary Tables

| **Feature** | **`gprofiler2`** | **`homologene`** | **`babelgene`** |
|---|---|---|---|
| Reference organisms | 700+ | 20+ | 19 (cannot convert between pairs of non-human species) |
| Gene mappings | More comprehensive | Less comprehensive | More comprehensive |
| Updates | Frequent | Less frequent | Less frequent |
| Orthology databases | Ensembl, HomoloGene, WormBase | HomoloGene | HGNC Comparison of Orthology Predictions (HCOP), including eggNOG, Ensembl Compara, HGNC, HomoloGene, InParanoid, NCBI Gene Orthology, OMA, OrthoDB, OrthoMCL, Panther, PhylomeDB, TreeFam, and ZFIN |
| Data location | Remote | Local | Local |
| Internet connection | Required | Not required | Not required |
| Speed | Slower | Faster | Medium |

**Supplementary Table S1. Comparison of orthology inference backends supported by `orthogene`.** Backends differ in organism coverage, data provenance, update frequency, and computational performance, allowing users to select an appropriate strategy based on analytical context.

**Supplementary Table S2. Gene orthology mapping options.** `orthogene::convert_orthologs` provides a variety of built-in strategies for handling gene orthology mappings that are not simply one-to-one (e.g. many-to-one, one-to-many, many-to-many). This simplifies and ensures consistency in data processing. These strategies can be employed by supplying either the full strategy name (*Option* column), the abbreviated name (*Abbreviation*), or a shorthand numeric ID for convenience (*Code*).

| Option | Abbreviation | Code | Description |
|---|---|---|---|
| drop_both_species | dbs | 1 | Drop genes that have duplicate mappings in either the input_species or output_species (DEFAULT). |
| drop_input_species | dis | 2 | Only drop genes that have duplicate mappings in the input_species. |
| drop_output_species | dos | 3 | Only drop genes that have duplicate mappings in the output_species. |
| keep_both_species | kbs | 4 | Keep all genes regardless of whether they have duplicate mappings in either species. |
| keep_popular | kp | 5 | Return only the most "popular" interspecies ortholog mappings. This procedure tends to yield a greater number of returned genes but at the cost of many of them not being true biological 1:1 orthologs. |
| sum / mean / median / min / max | – | – | When `gene_df` is a matrix and `gene_output="rownames"`, these options will aggregate many-to-one gene mappings (input_species-to-output_species) after dropping any duplicate genes in the output_species. |

#### 2 Supplementary Figures

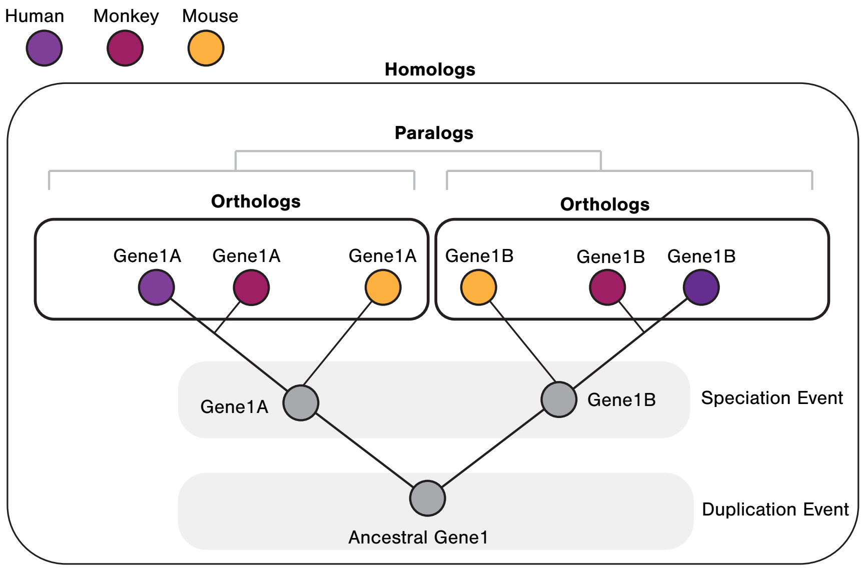


**Supplementary Fig. S1. Diagrammatic explanation of the differences between homologs, orthologs and paralogs.** *Homologs*: A broad supercategory of genes that are evolutionarily derived from a common ancestral gene. *Orthologs*: A subcategory of homologs derived from a single ancestral gene in the last common ancestor. *Paralogs*: A subcategory of homologs derived via a duplication event.

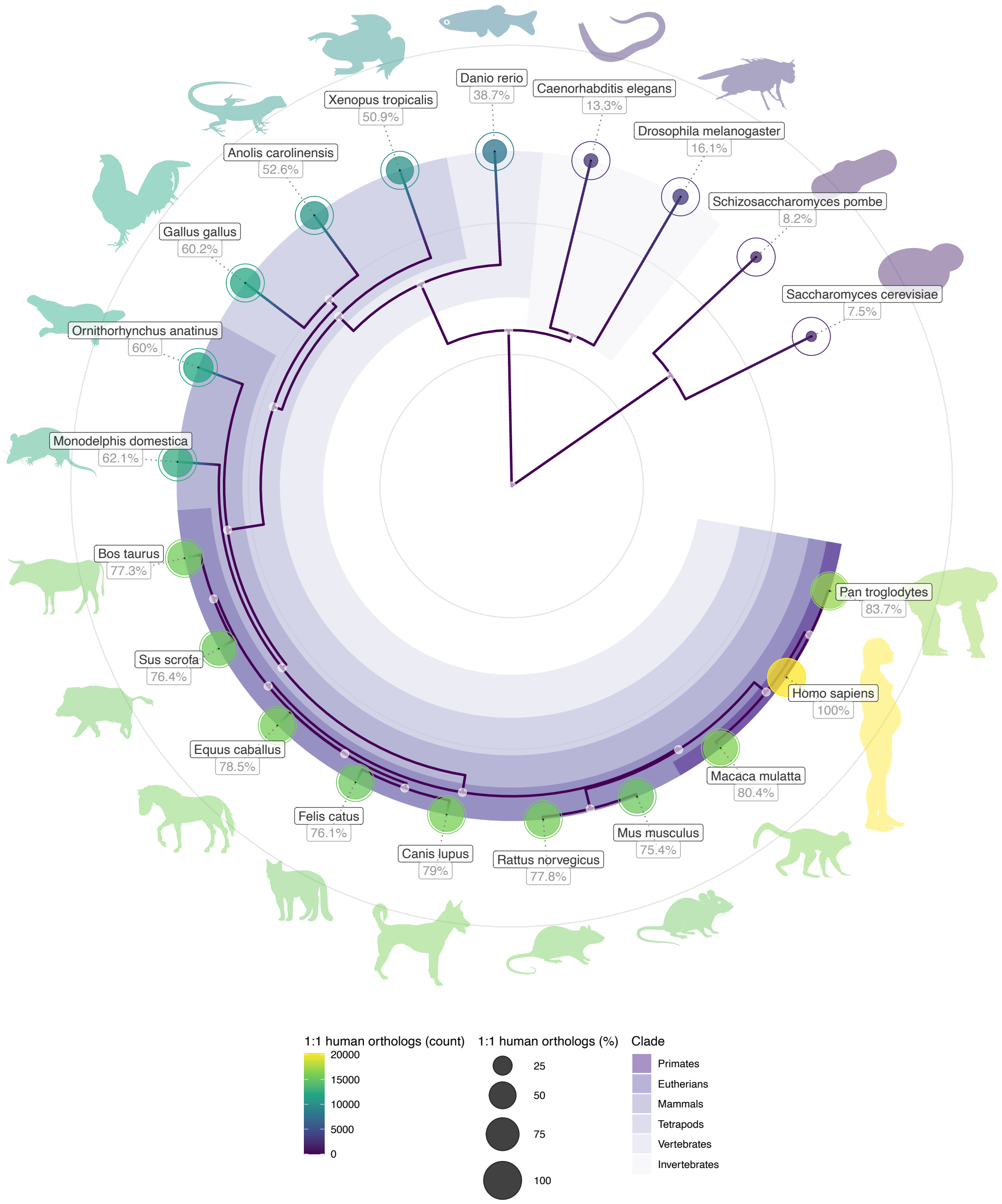


**Supplementary Fig. S2. Phylogenetic tree of 20 different organisms colored by the number of one-to-one homologs they share with humans.** Silhouette color indicates the number of one-to-one orthologs that species shares with humans. This plot was generated using *orthogene::plot_orthotree(method="babelgene")*. The percentage of human genes that have one-to-one homologs with each species is shown under the species scientific name label, and represented as the size of the inner dot within each species' respective circle.

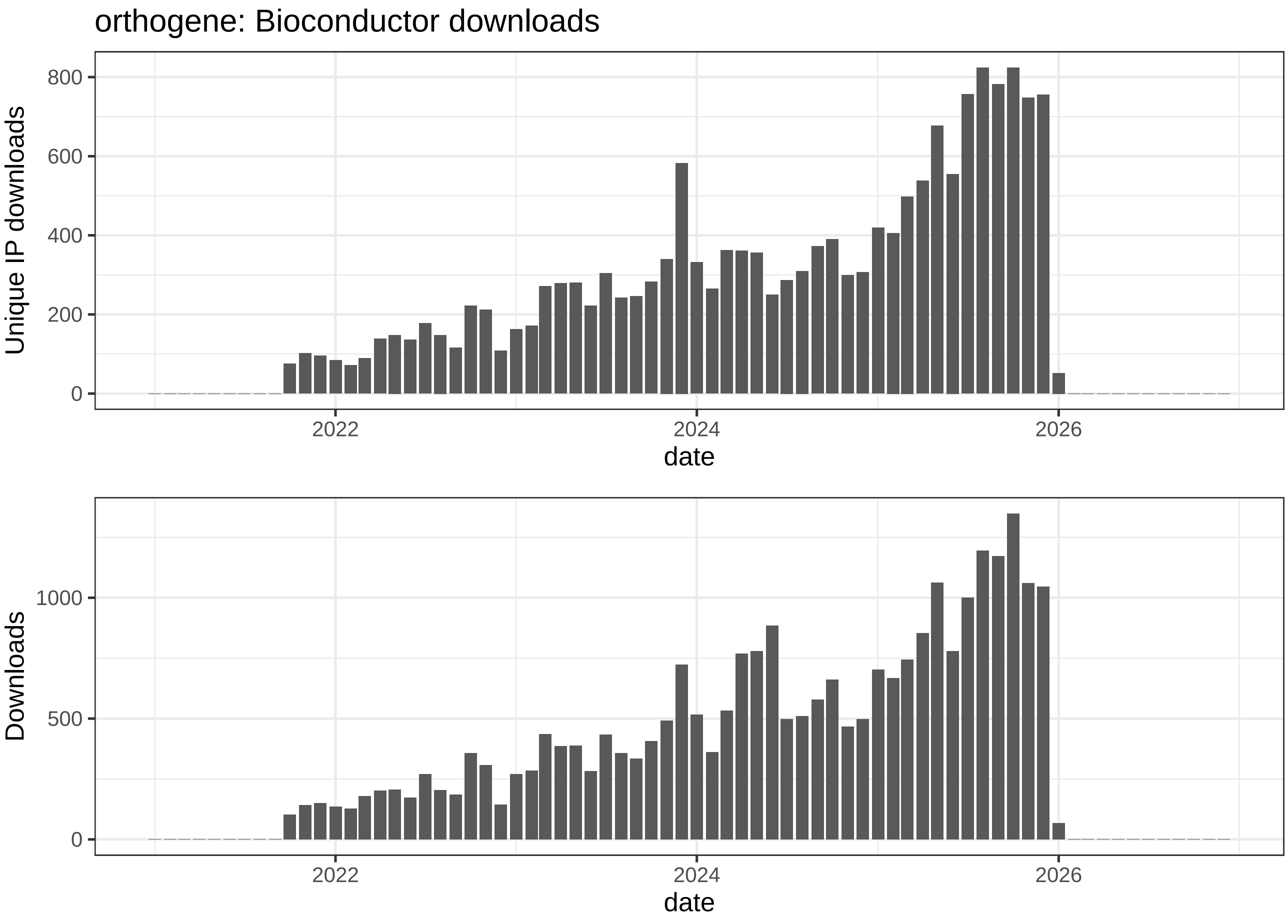


**Supplementary Fig. S3. `orthogene` has shown considerable uptake by the R community.** The number of unique IP addresses that downloaded `orthogene` each month since its release (*top*) and the total number of downloads each month (*bottom*).

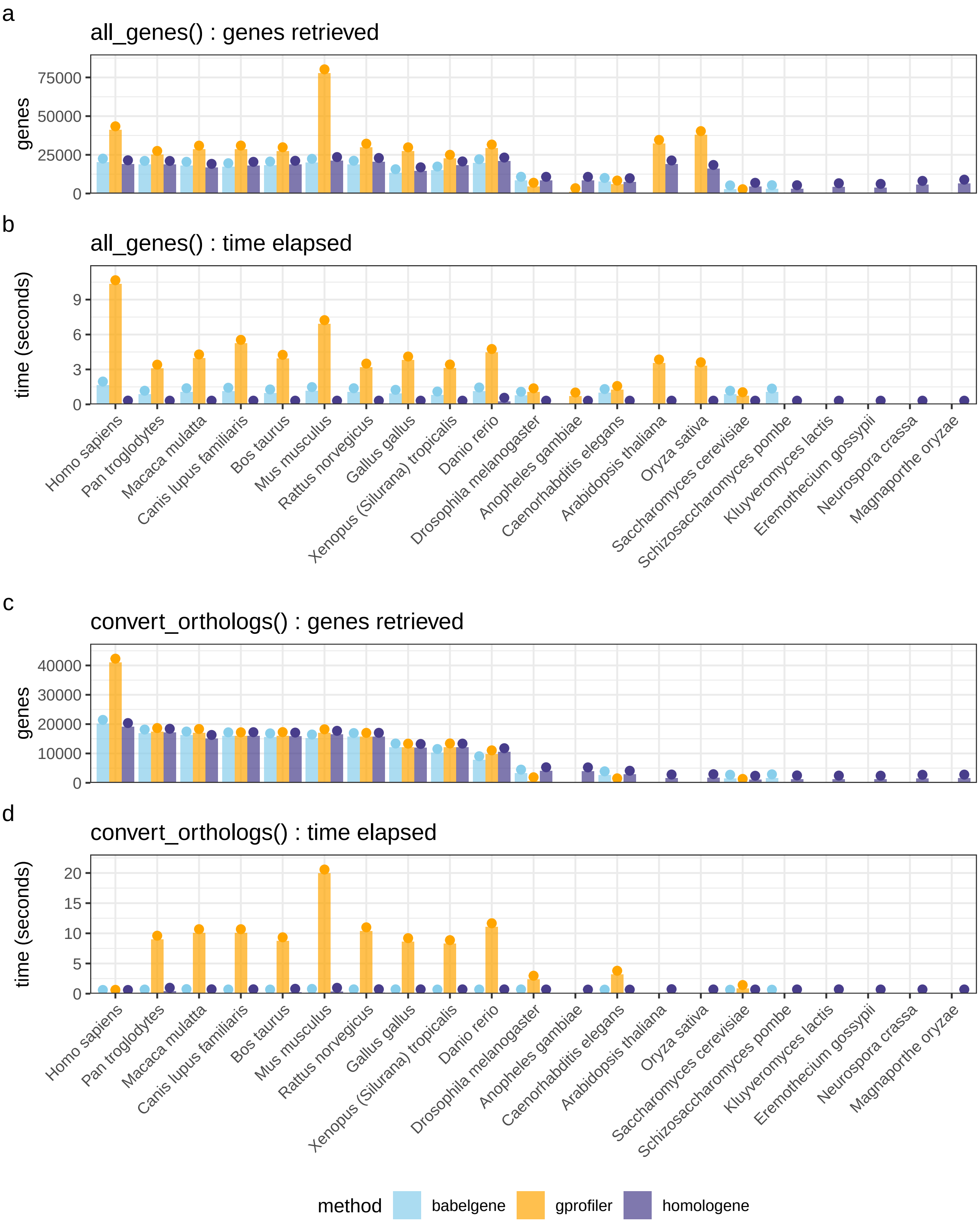


**Supplementary Fig. S4.** **Homology conversion backends show considerable variability in performance across model organisms. a**, The number of genes retrieved using `orthogene::all_genes()` with each backend method (`babelgene`, `gprofiler`, `homologene`). Missing dots above distinguish missing values (e.g. due to a species not being available with a certain method) from values close to or at 0. **b**, The time (seconds) it took for each method to run `orthogene::all_genes()`. **c**, The number of genes returned after running `orthogene::convert_orthologs()` using all human genes (from `orthogene::all_genes()` run with the same backend method) as input. **d**, Run time in seconds for each `orthogene::convert_orthologs()` function call.
